# Distinct labeling of host and viral genomes highlights redistribution of nuclear proteins during HSV-1 infection

**DOI:** 10.64898/2026.10.30.763179

**Authors:** Alison E. Lloyd, Jessica E. Packard, Sabrina L. Rutan, Jill A. Dembowski

**Affiliations:** Department of Biological Sciences, Duquesne University, Pittsburgh, PA, USA; Department of Pediatrics, University of Pittsburgh School of Medicine, Pittsburgh, PA, USA

## Abstract

Herpes simplex virus type 1 (HSV-1) coordinates viral and cellular proteins during infection. While proteins associated with HSV-1 genomes have been identified, changes in proteins associated with the host genome during infection remain poorly understood. We adapted accelerated native isolation of proteins on nascent DNA (aniPOND) to label and purify cellular and viral DNA then identify associated proteins by mass spectrometry. Using this approach, we confirm observations that cellular transcription, DNA replication, and repair proteins are enriched on viral genomes after the onset of viral DNA replication. We assessed how protein interactions with cellular DNA change during infection relative to protein abundance. Targets of the viral E3 ubiquitin ligase, ICP0, are depleted from the nucleus and cellular genome during infection. Transcription regulatory factors, Pol II, and the proteasome are depleted from the host genome, while association with DNA damage response and nuclear pore proteins increases. We also provide evidence for HSV-1 disruption of ribosome biogenesis. Results demonstrate reorganization of the nucleus during infection and validate this technique to identify changes to host genome-associated factors during infection.

**Author Summary:** Viruses remodel host cells to create an environment conducive to infection. Herpesviruses are known to alter host protein levels and localization to manipulate cellular processes and pathways. In this study, we explored how HSV-1 infection remodels host nuclei through changes to nuclear protein levels and association of proteins with cellular DNA using a novel approach to study host genome interactions during infection. Results provide a comprehensive view of how HSV-1 dramatically remodels host chromatin and nuclear processes and establishes a method that can be used to explore how nuclei are remodeled during infection with other DNA and RNA viruses.

## Introduction

Herpes simplex virus type 1 (HSV-1) is a prominent human pathogen, characterized clinically by lesions at the site of epithelial infection and the ability to establish latency and reactivate in sensory neurons. Like many other DNA viruses, HSV-1 has refined mechanisms for remodeling host nuclei and utilizing host machinery to drive viral processes. Understanding how HSV-1 remodels host cells is crucial for further elucidating the viral life cycle and infection outcome.

HSV-1 enters nuclei through nuclear pores as an approximately 152-kilobase-pair linear genome. Upon nuclear entry, HSV-1 gene expression occurs through a temporal cascade in which immediate early viral gene expression is driven by viral transcription factor and tegument protein, VP16 (Batterson & Roizman, 1983; Campbell *et al*, 1984). Two immediate early proteins, ICP4 and ICP0, are considered “master regulators” of HSV-1 infection and extensively remodel host nuclear architecture and processes to promote infection. ICP4 serves as the major viral transcription factor that promotes the recruitment of host transcription initiation factors and RNA polymerase II (Pol II) to drive expression of early and late viral genes (DeLuca *et al*, 1985; Dixon & Schaffer, 1980; Grondin & DeLuca, 2000; Watson & Clements, 1980). ICP4 preferentially binds to viral genomes that lack cellular nucleosomes, promoting Pol II recruitment to viral genes and contributing to host transcription shutoff (Dremel & DeLuca, 2019). ICP0 is an E3 ubiquitin-ligase that targets cellular viral restriction factors for proteasomal degradation (Boutell *et al*, 2002; Everett *et al*, 2006; Maul *et al*, 1993), including components of promyelocytic leukemia protein (PML) nuclear bodies such as PML, SP100, and other SUMOylated proteins (Everett *et al*, 1998; Rodriguez *et al*, 2020). ICP0 also targets key components of the cellular DNA damage response (DDR) to positively influence infection (Lilley *et al*, 2011; Parkinson *et al*, 1999).

Viral replication compartments are hallmarks of HSV-1 infection, containing viral genomes undergoing DNA replication, recombination, gene expression, and virion assembly (Kobiler & Weitzman, 2019; Packard & Dembowski, 2021). As viral replication compartments expand, host chromatin adopts a highly compacted, transcriptionally inactive state at the nuclear periphery (Aho *et al*, 2019; Gonzalez-Almela *et al*, 2025). Replication compartments are largely nucleosome-free during lytic replication (McSwiggen *et al*, 2019; Nosek *et al*, 2026), enabling accumulation of viral and host factors involved in transcription, DNA replication, DNA repair, chromatin remodeling, and virion assembly (Dembowski & DeLuca, 2015, 2018). Despite chromatin compaction, infection results in increased nuclear fluidity, which is mediated by ICP4 and enables nuclear condensate formation and viral replication compartment maturation (Herzog *et al*, 2026).

Further evidence of HSV-1 compartmentalization of the host nucleus is the formation of virus-induced chaperone-enriched (VICE) domains. These domains form at the periphery of replication compartments and sequester host chaperone protein heat shock cognate protein 70 (Hsc70) (Burch & Weller, 2004). VICE domains also contain components of the 20S proteasome and exhibit proteolytic activity (Livingston *et al*, 2009).

Taken together, viral infection causes dramatic reorganization of host nuclear compartments to promote viral replication. While extensive work has characterized proteins associated with viral genomes (Dembowski & DeLuca, 2015, 2018; Dembowski *et al*, 2017; Reyes *et al*, 2017), comparatively little is known about how the cellular genome-associated proteome is altered during infection. This study aimed to determine how protein association with host chromatin changes during HSV-1 infection. To do this, we adapted accelerated native isolation of proteins on nascent DNA (aniPOND) (Dembowski & DeLuca, 2015, 2017; Leung *et al*, 2013; Reyes *et al*., 2017) to label and purify cellular DNA and identify associated proteins in infected and uninfected cells. Through a combination of cellular and viral aniPOND, nuclear protein purification, and imaging, we assessed how protein interactions with cellular DNA change during infection compared to uninfected cells, relative to changes in nuclear protein abundance and protein association with viral DNA. Collectively, results from this study provide a model describing how nuclear proteins and compartments are highly reorganized during HSV-1 infection.

## Results

### An approach to selectively label cellular DNA for investigation during infection

We previously used ethynyl-modified nucleosides to label replicating viral DNA to study protein association with viral replication compartments (Dembowski & DeLuca, 2015) or forks (Dembowski *et al*., 2017), and have prelabeled viral DNA within virions to investigate the temporal association of proteins with infecting viral genomes (Dembowski & DeLuca, 2018). Here, we sought to selectively label cellular DNA to investigate how infection alters protein association with host genomes. To exclusively label cellular DNA and not viral DNA, MRC-5 cells were plated at a low density and 5-ethynyl-2‘-deoxycytidine (EdC) was added to the growth medium to be incorporated into replicating DNA for 48 hours. Then, EdC was washed out and cells were incubated for another 24 hours without EdC, followed by mock or HSV-1 infection (Fig 1A). For comparison, to exclusively label viral DNA and not cellular DNA, MRC-5 cells were plated and grown under the same conditions in the absence of EdC, followed by infection and replication compartment labeling with EdC from 4 to 6 hours post infection (hpi). We previously confirmed that under these conditions viral DNA and not cellular DNA is labeled with EdC (Dembowski & DeLuca, 2015; Dembowski *et al*., 2017). For all conditions, after 6 hours of mock or HSV-1 infection, EdC-labeled viral or cellular DNA was attached to biotin for aniPOND and proteomic analysis or a fluorophore for imaging (Fig 1B). As a negative control for aniPOND, cells were incubated in the absence of EdC to identify background binding.

**Figure 1.**
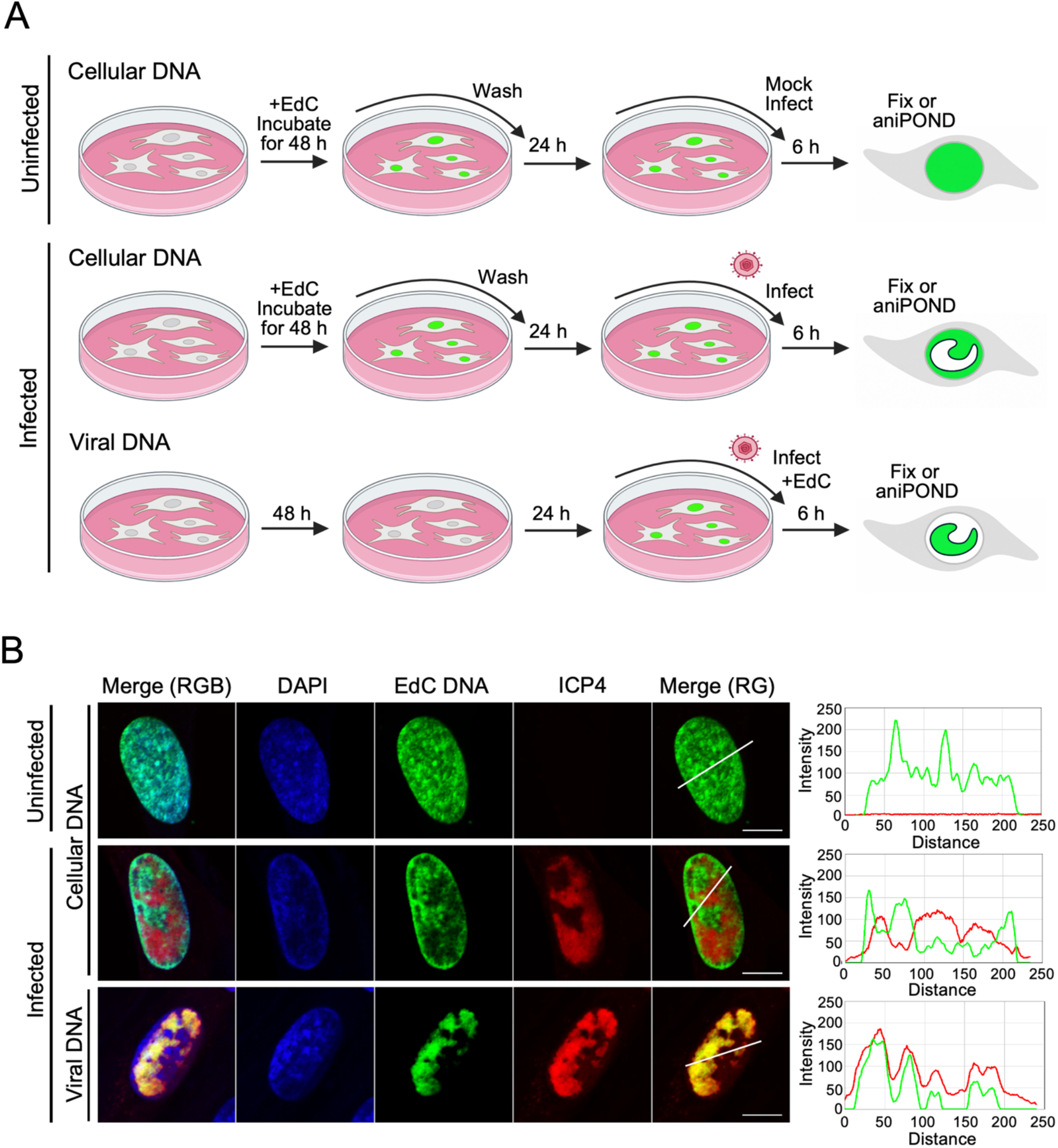
Selective labeling of cellular and HSV-1 DNA with EdC. A) Schematic workflow of selective EdC labeling in MRC-5 cells. Cellular DNA was labeled with 25 μM EdC and cells were either mock-infected or infected with KOS at an MOI of 10 PFU/cell for 6 hours. To label viral DNA, cells were grown under the same conditions, infected, and EdC was added from 4 to 6 hpi. Cells were then fixed for immunofluorescence or processed for aniPOND. Image generated in BioRender. B) Images of EdC-labeled cellular and viral DNA. Infection and EdC labeling were carried out as indicated in (A). Cellular DNA was stained with DAPI (blue), EdC-labeled DNA was tagged via click chemistry to Alexa Fluor 488 (green), and viral protein ICP4 was labeled using indirect immunofluorescence (red). Scale bars, 10 μm. Trace files, generated using ImageJ, show the relative intensity of tagged EdC-labeled DNA (green) and ICP4 immunofluorescence (red) along the line shown in the Merge (RG) panel.

As prolonged exposure to nucleoside analogs can affect cell viability, a CellTiter-Glo Luminescent Cell Viability Assay was conducted to assess the cytotoxic effects of adding EdC to replicating cellular DNA for 48 hours (Fig S1). EdC was not cytotoxic at or below 50 μM. Thus, 25 μΜ EdC was used for all labeling.

To validate the selective labeling of cellular and viral DNA during HSV-1 infection, EdC-labeled genomes were visualized by click chemistry and immunofluorescence microscopy (Fig 1B). In uninfected cells, cellular DNA fills the nucleus with regions of more intense signal, likely corresponding to cellular heterochromatin. In infected cells, cellular DNA appears more marginalized to the nuclear periphery, with markedly less intense staining in regions containing viral replication compartments, which were labeled by ICP4 staining. Distinct patterns of nuclear staining were observed between labeled cellular DNA and viral ICP4. Conversely, labeled viral DNA and ICP4 displayed a clear overlap. This visualization confirms the use of EdC labeling and click chemistry to identify distinct populations of cellular and viral DNA during infection.

### Proteins that regulate distinct processes are differentially enriched on viral compared to cellular DNA during infection

Having established that host and viral DNA can be selectively labeled with EdC (Fig 1), we utilized aniPOND coupled with mass spectrometry (MS) to identify and compare the proteomes associated with viral and host DNA at 6 hpi. The normalized spectral abundance factor (NSAF) of viral and host DNA-associated proteins was calculated and compared to determine the relative enrichment of proteins on viral or host DNA (Fig 2A, Table S1). The right side of the volcano plot reveals proteins enriched on viral DNA at 6 hpi by greater than 2-fold compared to cellular DNA (Fig 2A, blue). Significantly enriched proteins include transcription regulatory factors ICP4 and the Mediator complex. Also enriched on viral DNA are DNA replication and repair factors, including viral origin-binding protein UL9, proliferating cell nuclear antigen (PCNA), and mismatch repair protein MutS homolog 2 (MSH2). On the other hand, proteins enriched by at least 2-fold on host DNA are shown on the left (red). Statistically significant proteins include 60S ribosome assembly factors nucleolar protein 56 (NOP56) and MRT4 homolog (MRTO4), ribosomal protein RPL29, chromatin regulatory proteins methy-CpG binding protein (MECP2) and chromatin target of PRMT1 (CHTOP), histone protein HIST1H1C, and nuclear pore protein NUP153. These results demonstrate that HSV-1 infection results in statistically significant differences in protein recruitment to viral or cellular genomes.

**Figure 2.**
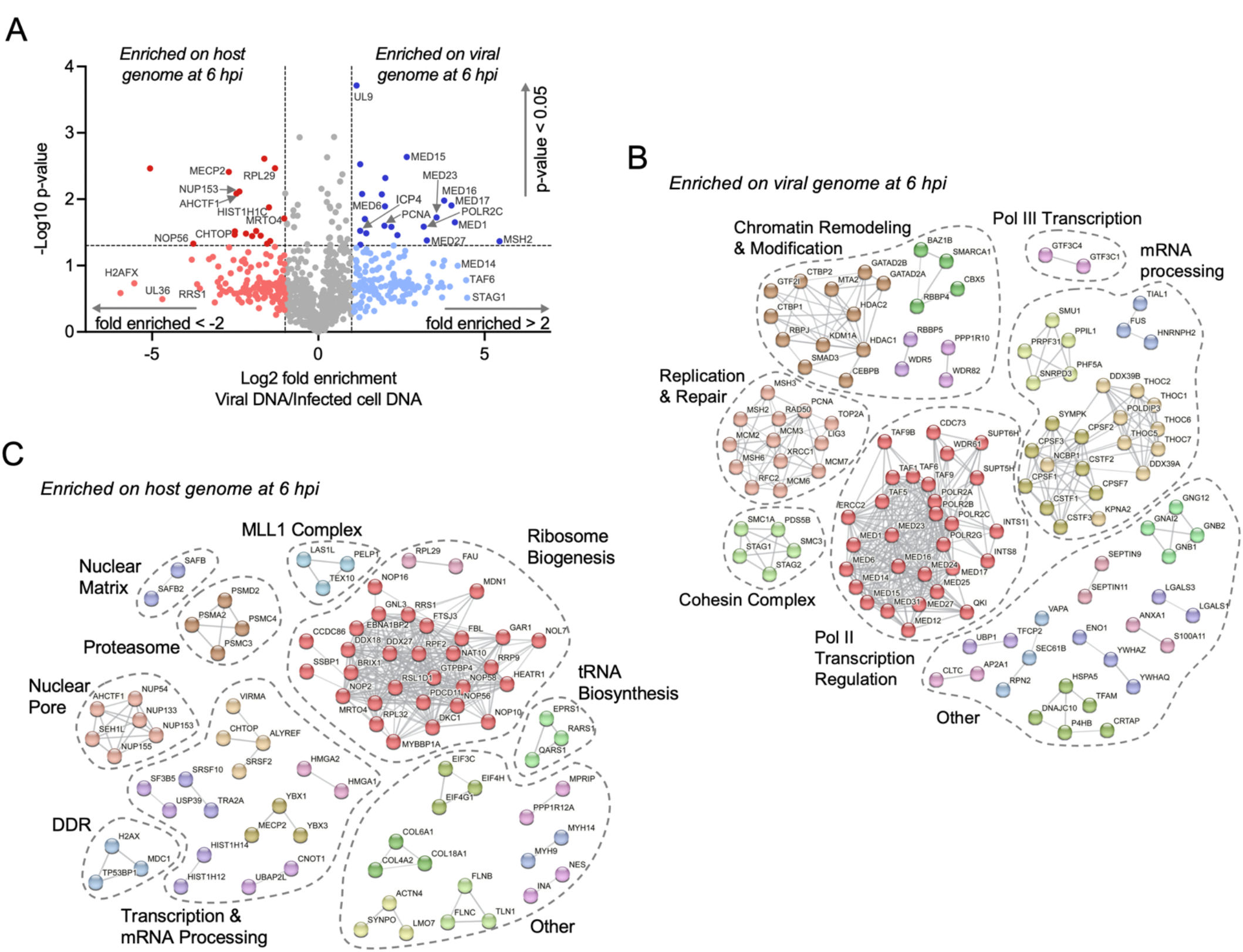
Relative association of proteins with cellular and viral genomes during HSV-1 infection. A) Volcano plot of host and viral proteins bound to EdC-labeled viral DNA compared to proteins bound to EdC-labeled cellular DNA. All proteins were isolated from EdC-labeled DNA at 6 hpi by aniPOND, as described in Figure 1A (infected conditions). The x-axis represents the log 2-fold difference in protein enrichment, and the y-axis displays statistical significance as-log_10_ p-value. Fold difference and p-value were calculated using log-transformed NSAF values. Horizontal and vertical lines represent significance thresholds at p=0.05 and a two-fold difference. Each data point represents an individual protein detected in three replicate experiments and is color-coded based on significance and fold difference in enrichment on viral versus cellular DNA during infection. Proteins in red were considered enriched on the host genome and proteins in blue were considered enriched on the viral genome. Proteins with no significant difference in enrichment or with less than a two-fold difference in enrichment on viral versus cellular DNA are shown in grey. A complete list can be found in Table S1. B-C) STRING predicted interaction network between proteins enriched on viral (B) or cellular (C) genomes at 6 hpi. Proteins that were not clustered by STRING analysis but considered enriched on viral or cellular genomes during infection are listed in Figure S2.

To further characterize viral and host genome-associated cellular proteins during infection, STRING protein interaction networks were generated to identify groups of proteins that are relatively more enriched on viral (Fig 2B) or cellular (Fig 2C) DNA. Consistent with previous studies (Dembowski & DeLuca, 2015), proteins enriched on viral genomes at 6 hpi included factors involved in transcription regulation, DNA replication and repair, chromatin remodeling and modification, and mRNA processing (Fig 2B, S2A). Transcription regulatory complexes include transcription factor IID (TFIID), the Mediator complex, the Integrator complex, and Pol II subunits. In addition, cellular replication proteins PCNA, DNA ligase 3 (LIG3), and topoisomerase 2A (TOP2A), as well as mismatch repair (MMR) proteins and double-strand break repair protein RAD50 were relatively more enriched on viral than cellular DNA during infection. The cohesin complex was also enriched on viral DNA, which in uninfected cells is known to hold replicated sister chromatids together and contribute to gene expression and DNA repair. Taken together, comparing proteins relatively more enriched on viral than cellular DNA highlights the robust expression and replication of viral DNA at 6 hpi.

In contrast, proteins enriched on cellular genomes during infection are largely associated with ribosome biogenesis, transcription regulation, mRNA processing, the nuclear pore, and nuclear matrix (Fig 2C, S2B). Although select transcription and mRNA processing factors were relatively more enriched on cellular compared to viral DNA during infection, these were far less clustered and did not include the core complexes required for expression of all Pol II-regulated genes as was found associated with viral DNA. Most notably enriched on host DNA were factors involved in ribosome biogenesis. Most of these proteins contribute to the assembly and maturation of 60S ribosomal subunits, including BRIX1, NOP proteins, midasin AAA ATPase 1 (MDN1), and DEAD-box RNA helicases (DDX). As is the case for mRNA processing, ribosome assembly factors were likely captured by aniPOND because they participate in pre-rRNA processing events that occur while the rRNA is still attached to the rDNA locus on the cellular genome. We also observed enrichment of select mediators and markers of the DDR, including histone H2AX, mediator of DNA damage checkpoint 1 (MDC1), and tumor protein p53 binding protein 1 (TP53BP1). These are key components of DNA damage detection and signaling pathways. These findings reveal that although Pol II-mediated transcription, DNA replication, and repair mostly occur on viral DNA during infection, distinct transcription regulatory proteins, ribosome assembly factors, and proteins that facilitate the DDR are recruited to host genomes.

### Viral infection results in a robust shift in nuclear protein levels and association with cellular chromatin

To further investigate nuclear reorganization during HSV-1 infection, we compared total isolated nuclear proteins with those bound to host DNA. The first part of this multi-step comparison involved isolation and identification of total nuclear proteins from uninfected and infected cells at 6 hpi. Statistical analyses revealed subsets of proteins that increased or decreased in relative abundance upon infection, as well as a large set of proteins with minimal change (Fig 3A, 5A, Table S2). Next, to determine how HSV-1 alters protein association specifically with host chromatin, aniPOND was performed on EdC-labeled cellular DNA in uninfected and infected cells at 6 hpi. While a large subset of proteins showed minimal change between conditions, select proteins were found to be enriched on or depleted from host DNA during infection (Fig 3B, 5B, Table S1).

**Figure 3.**
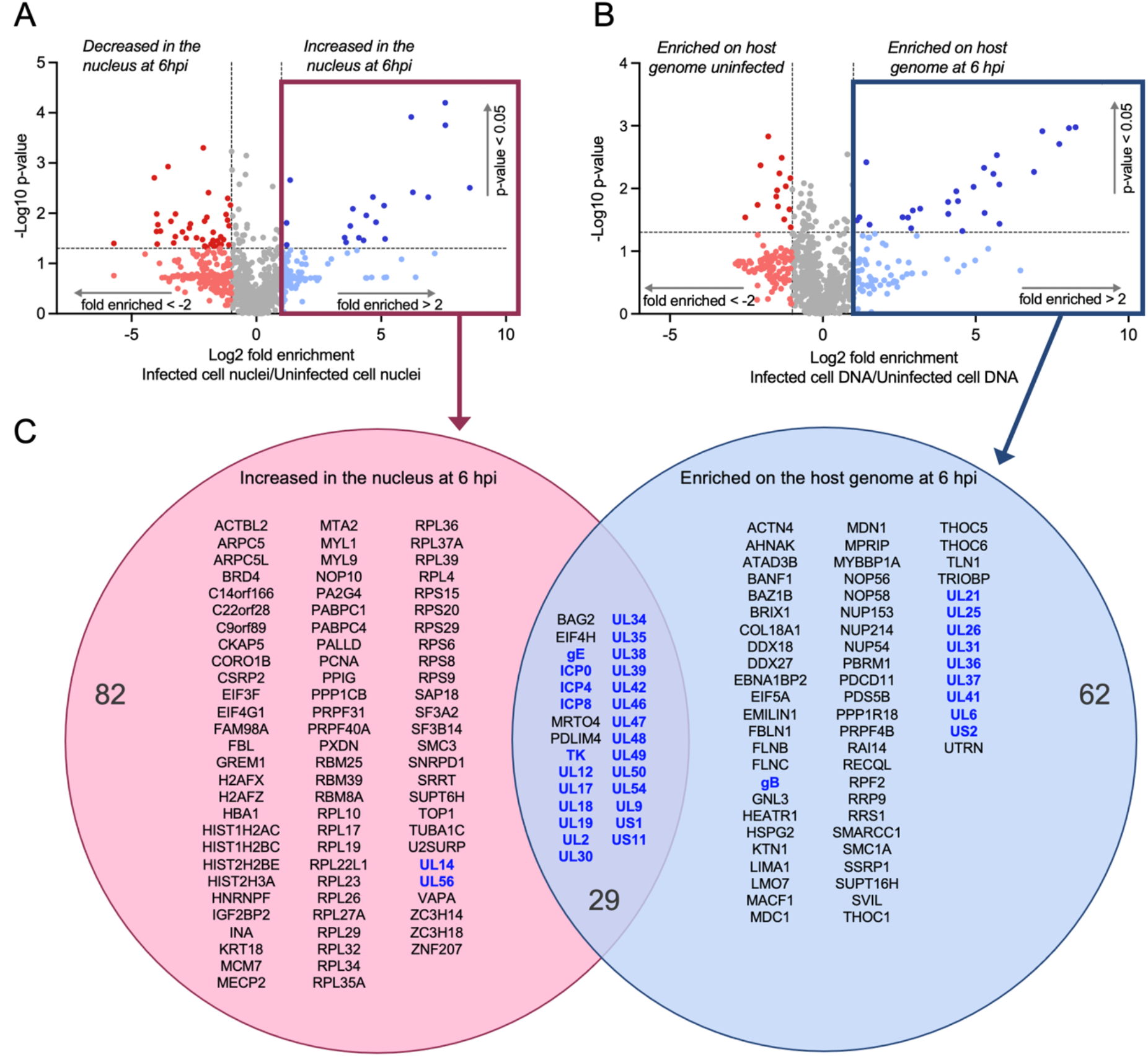
Comparison of proteins enriched in the nucleus vs. on cellular genomes during infection. A) Volcano plot of cellular and viral proteins isolated from infected cell nuclei compared to mock-infected cell nuclei at 6 hpi. B) Volcano plot of cellular and viral proteins bound to EdC-labelled cellular DNA in infected cells, compared to proteins bound to EdC-labelled cellular DNA in mock-infected cells. For both plots, the x-axis represents log2 fold difference (infected/uninfected), and the y-axis displays statistical significance as-log_10_ p-value. Paired t-tests of log-transformed NSAF values from 3 biological replicates were used to calculate p-values. Horizontal and vertical lines represent significance thresholds at p=0.05 and a two-fold change. Each data point represents an individual protein detected in three replicate experiments and is color-coded based on significance and fold difference in enrichment. Proteins in blue were considered increased in the nucleus (A) or on the host genome (B) during infection, and proteins in red were considered decreased in the nucleus (A) or on the host genome (B) during infection. Proteins with no significant difference in enrichment or with less than a two-fold difference are shown in grey. A complete list can be found in Table S2 (A) or Table S1 (B). C) Venn diagram of proteins identified by nuclear isolation or aniPOND to be increased in the nucleus (pink) and/or enriched on cellular DNA (blue) at 6 hpi. HSV-1 proteins are shown in blue.

Utilizing these two datasets, we examined if proteins that increased in the nucleus were also enriched on host chromatin during infection. Comparison of proteins meeting the 2-fold enrichment threshold identified 29 proteins to be increased in the nucleus and on host chromatin during infection, while 82 and 62 proteins were uniquely increased in the nucleus or on host DNA, respectively (Fig 3C). The overlap between proteins that increase in the nucleus and on host chromatin largely includes viral proteins; although several proteins relating to distinct cellular functions were also identified, including the BAG family molecular chaperone 2 (BAG2), eukaryotic translation initiation factor 4H (EIF4H), tumor suppressor PDZ and LIM domain 4 (PDLIM4), and mRNA turnover protein 4 homolog (MTRO4), which has been implicated in 60S ribosome assembly and mRNA turnover.

Proteins that uniquely increased in the nucleus or on host DNA were then subject to STRING functional analysis. Proteins that increased in the nucleus but not on cellular DNA form interaction networks associated with mRNA processing, chromatin organization, and ribosomal proteins, as well as select proteins involved in DNA replication, tRNA biosynthesis, and ribosome biogenesis (Fig 4A). Although ribosomal protein levels increase in the nucleus, they do not increase on the host or viral genome during infection. This is consistent with studies that have shown that ribosomal proteins localize to VICE domains during infection (Metzger *et al*, 2025). Proteins that increased on the host genome, but not in the nucleus upon infection primarily included factors involved in ribosome biogenesis and chromatin remodeling, as well as the DDR protein MDC1 and nuclear pore, mRNA export, and cohesion proteins (Fig 4B). Note that some viral proteins increased on the host genome but not in the nucleus. Logically, these proteins increase in the nucleus during infection indicating that aniPOND-MS is more sensitive to detect viral proteins than nuclear MS.

**Figure 4.**
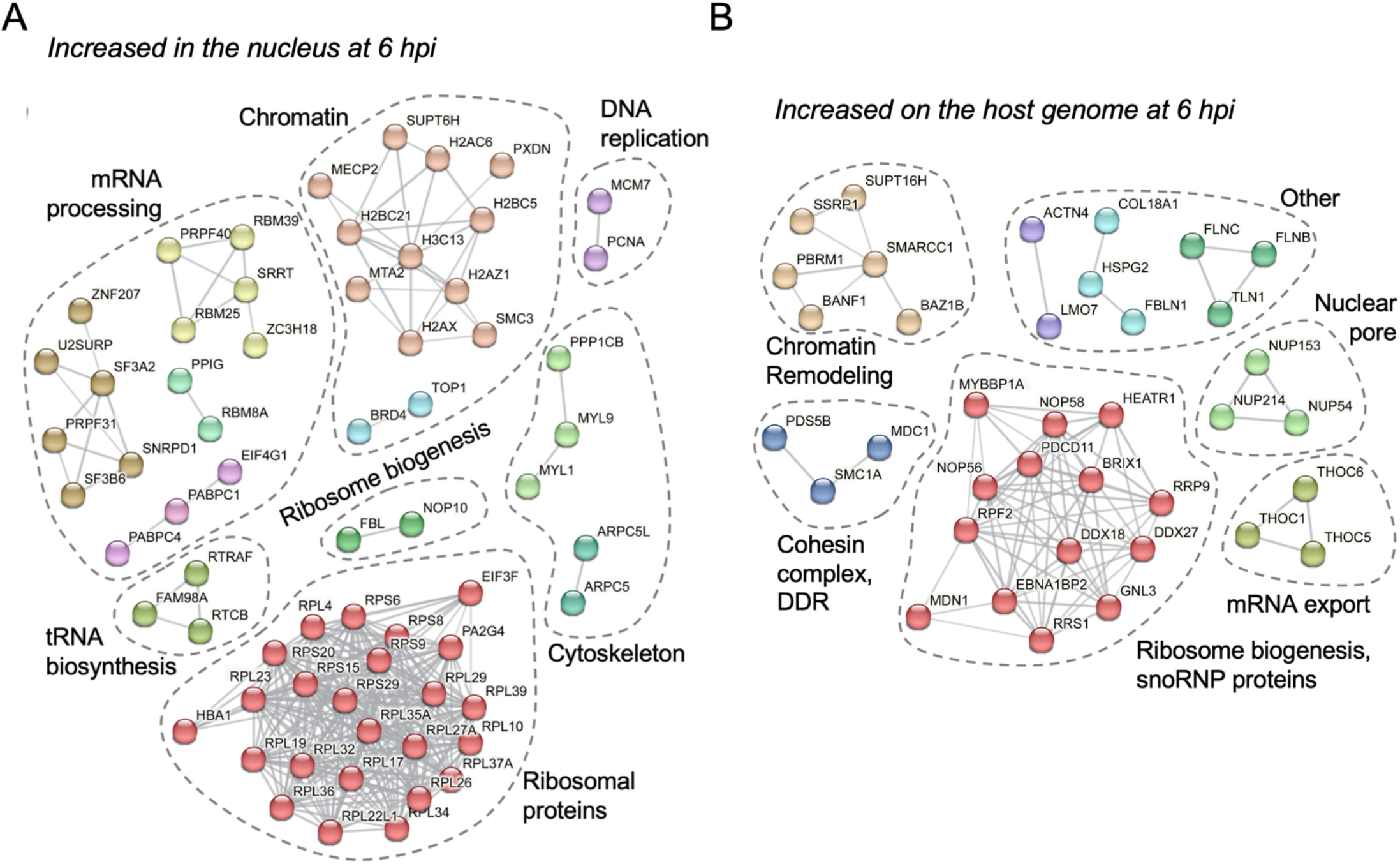
STRING analysis of proteins that increase in the nucleus or on the host genome at 6 hpi. A) STRING interaction network of proteins that increased in infected cell nuclei but not on cellular DNA at 6 hpi (Figure 3C, pink). B) STRING interaction network of proteins that increased association with cellular DNA but not in the nucleus at 6 hpi (Figure 3C, blue). Proteins are color-coded according to STRING cluster categories. Proteins that were unmapped due to lower confidence interactions can be found in the full list in Figure 3C.

Next, we compared the converse relationship, proteins that decreased in nuclear abundance to proteins depleted from host chromatin during infection (Fig 5). Comparison of proteins that decreased in the nucleus and on cellular genomes by more than 2-fold revealed 13 proteins simultaneously depleted in the nucleus and from host DNA (Fig 5C). Notably, this overlap included select DDR proteins, including DNA-dependent protein kinase (DNA-PK, PRKDC gene), a target of ICP0, and TP53BP1. Although TP53BP1 was found to be more enriched on cellular than viral DNA (Fig 2C), total protein levels in the nucleus and association with host chromatin decreased during infection (Fig 5C). Chaperonin proteins (CCT7, CCT8) and proteasomal components (PSMC1, PSMD1) were also found in this overlap. In addition, 209 proteins were identified to decrease in the nucleus during infection, but not on cellular DNA, and 91 proteins we identified to decrease association with cellular DNA, but not in the nucleus.

**Figure 5.**
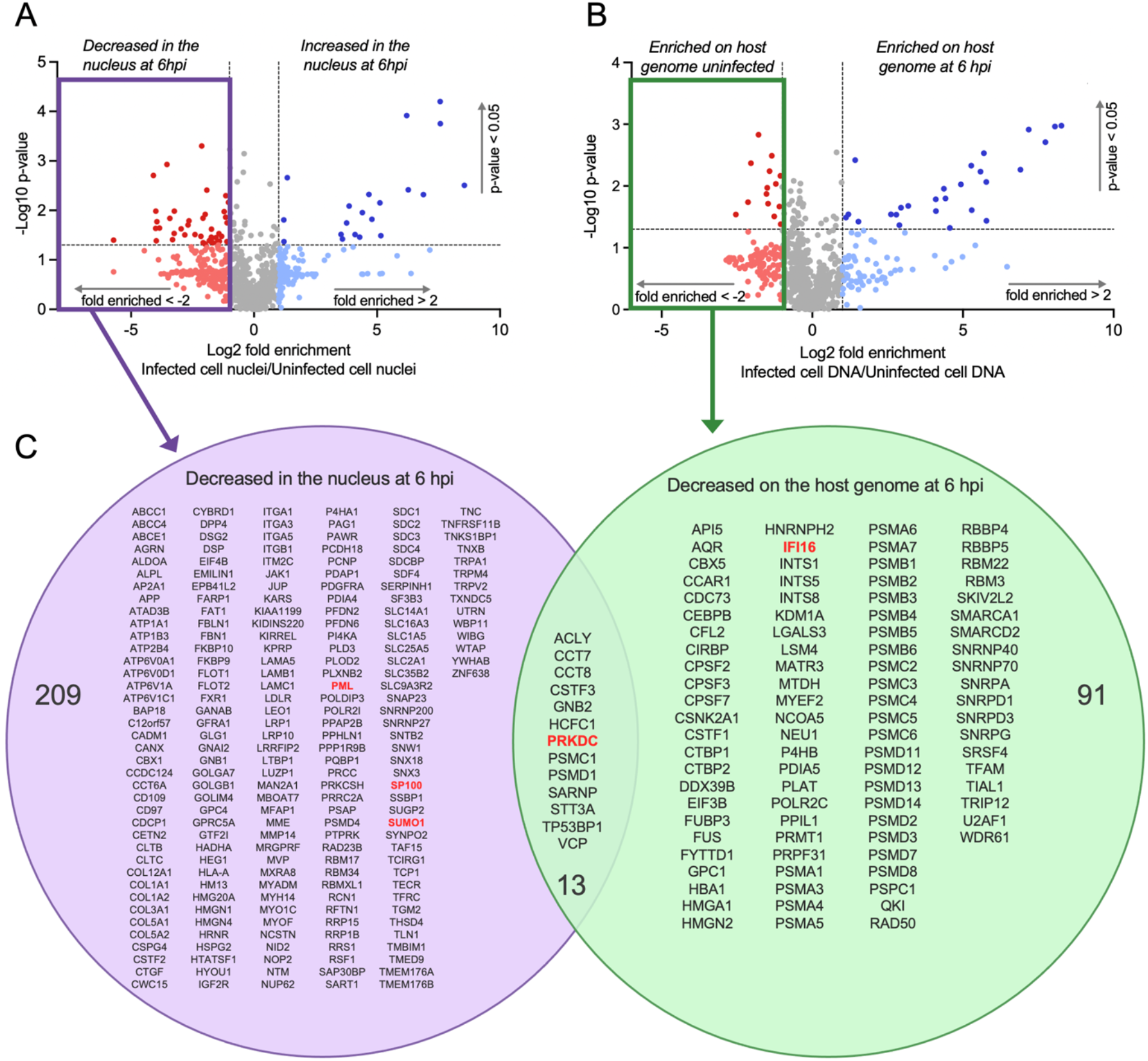
Comparison of proteins that decrease in the nucleus vs. on the cellular genome during infection. A.) Volcano plot of host and viral proteins isolated from infected cell nuclei compared to mock-infected cell nuclei at 6 hpi. B.) Volcano plot of host and viral proteins bound to EdC-labelled cellular DNA in infected cells compared to proteins bound to EdC-labelled cellular DNA in mock-infected cells. For both plots, the x-axis represents log2 fold difference (infected/uninfected), and the y-axis displays statistical significance as-log_10_ p-value. Paired t-tests of log-transformed NSAF values from 3 biological replicates were used to calculate p-values. Horizontal and vertical lines represent significance thresholds at p=0.05 and a two-fold change. Each data point represents an individual protein, which is color-coded based on significance and fold difference in enrichment. Proteins in blue were considered increased in the nucleus (A) or on the host genome (B) during infection, and proteins in red were considered decreased in the nucleus (A) or on the host genome (B) during infection. Proteins with no significant difference in enrichment or with less than a two-fold difference are shown in grey. A complete list can be found in Table S2 (A) or Table S1 (B). C) Venn diagram comparing proteins identified by nuclear isolation or aniPOND to be decreased in the nucleus (purple) and/or on cellular DNA (green) at 6 hpi. ICP0 targets are shown in red.

Proteins uniquely depleted from the nucleus during infection formed a broad STRING interaction network representing diverse cellular processes (Fig 6A). One of the most notable functional groups depleted included PML nuclear body proteins (PML, SUMO1, SP100), which are targeted for proteasomal degradation by ICP0 (Everett *et al*., 1998; Rodriguez *et al*., 2020). In addition, mRNA processing factors, including spliceosome-associated proteins, decreased in the nucleus, perhaps contributing to the mechanism by which HSV-1 infection disrupts spliceosome function (Hardy & Sandri-Goldin, 1994). Select proteins involved in Pol II transcription regulation, ribosome biogenesis, DNA repair, signaling, the unfolded protein response, stress response, and the chaperonin complex also decreased. On the other hand, STRING functional analysis revealed a distinct collection of cellular proteins depleted from the host genome during infection (Fig 6B). Proteins that decreased association with the cellular genome but did not decrease in the nucleus include much of the proteasome and proteins and complexes that regulate Pol II-mediated transcription, cleavage and polyadenylation, RNA processing, and chromatin remodeling. As shown in Figure 2B, many of these proteins are re-localized to viral genomes and previous studies demonstrated that the proteasome is sequestered in VICE domains (Burch & Weller, 2004). Taken together, this study provides a comprehensive view of how host nuclear architecture is rearranged during HSV-1 infection.

**Figure 6.**
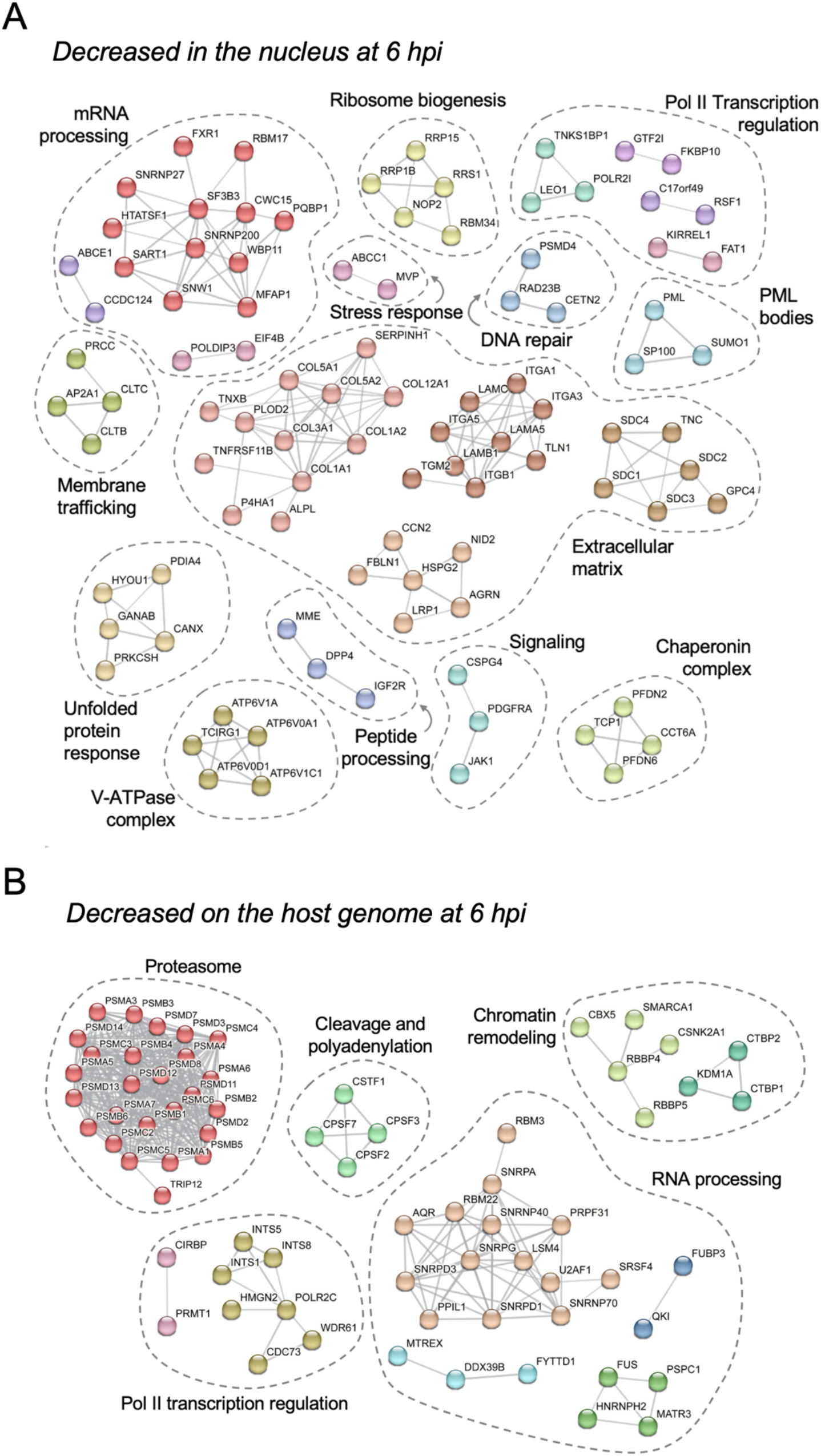
STRING analysis of proteins that decrease in the nucleus or are deficient on the host genome at 6 hpi. A) STRING interaction network of proteins that decreased in infected cell nuclei, but not on cellular DNA at 6 hpi (Figure 5C, purple). B) STRING interaction network of proteins that decreased association with cellular DNA but not in the nucleus at 6 hpi (Figure 5C, green). Proteins are color-coded according to STRING cluster categories. Proteins that were unmapped due to lower confidence interactions can be found in the full list in Figure 5C.

### Viral infection alters nucleolar architecture

Increased association of select ribosome assembly factors with host chromatin and ribosomal proteins in the nucleus during infection suggests a specific alteration in ribosome assembly. It was surprising that several ribosome biogenesis factors were enriched on cellular DNA during infection. However, we propose that the increased association was a result of tethering of stalled preribosomes with rDNA. We therefore used immunofluorescence imaging and click chemistry as in Figure 1B to determine how the localization of a nucleolar 60S ribosome assembly factor BRIX1 changes during infection (Fig 7A). In infected cells, BRIX1 localizes to a nucleolar structure that lacks viral DNA and host chromatin. This is consistent with the hypothesis that the increased association of nucleolar factors with cellular DNA during infection represents a ribosome assembly intermediate tethered to EdC-labeled cellular DNA. In addition, we consistently observed altered nucleolar structure with less rounded and defined nucleoli during infection compared to a more compacted appearance in uninfected cells (Fig. 7B).

**Figure 7.**
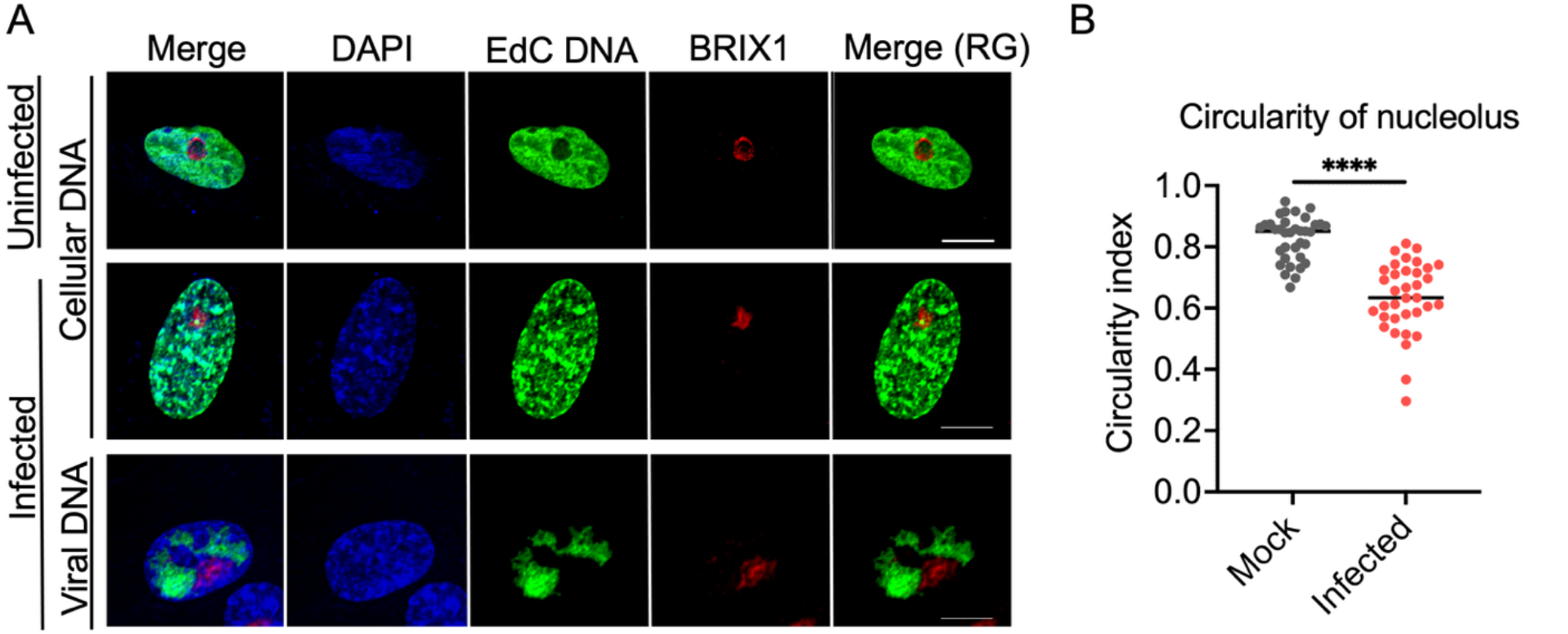
BRIX1 localization and nucleolar structure during HSV-1 infection. Infection and EdC labeling were carried out as indicated in Figure 1A. Cellular DNA was stained with DAPI (blue), EdC-labeled DNA was tagged via click chemistry to Alexa Fluor 488 (green), and ribosome assembly factor BRIX1 was labeled using indirect immunofluorescence (red). Scale bars, 10 μm. Trace files, generated using ImageJ, show the overlap between BRIX1 (red) and labeled cellular or viral DNA (green) along the line shown in the Merge (RG) panel. B) Circularity was measured using BRIX1 as a nucleoli marker where 1.0 is a perfect circle. An unpaired t-test with Welch’s correction was used to compare circularity of the nucleolus in the mock vs infected conditions (n.s. not significant, * p<0.0.05, ** p <0.01, *** p<0.001, **** p<0.0001).

## Discussion

Viral infection results in the robust remodeling of host cells and manipulation of cellular processes. For herpesviruses, nuclear reorganization is key to facilitating viral genome replication, gene expression, and the assembly and exit of nascent virions. A comprehensive analysis of how host genome interactions are modified by infection is essential to understanding this nuclear reorganization. Here, we utilized aniPOND to purify cellular DNA from infected and uninfected cells for proteomic analysis of cellular genome-associated proteins to determine how infection alters host chromatin. To exclusively label cellular DNA with EdC and avoid viral DNA contamination, we prelabeled replicating cellular DNA with EdC prior to HSV-1 infection, followed by purification and MS identification of genome-associated proteins after the onset of viral DNA replication. For comparison, we also identified proteins enriched on uninfected cell DNA and viral replication compartments and determined how nuclear protein levels change during infection. Results are supported by previous studies that have shown that infection results in cellular genome compaction and marginalization (Aho *et al*., 2019; Bosse *et al*, 2015), shut down of host DNA replication and gene expression (Dremel & DeLuca, 2019), depletion of ICP0 targets and disruption of PML nuclear bodies (Everett *et al*., 2006; Lilley *et al*., 2011; Maul *et al*., 1993; Mertens & Knipe, 2021; Rodriguez *et al*., 2020), formation of VICE domains containing the proteasome (Burch & Weller, 2004; Livingston *et al*., 2009), increased nuclear localization of ribosomal proteins and accumulation in VICE domains (Metzger *et al*., 2025), stalled ribosome biogenesis (Belin *et al*, 2010), and altered host DDR and DNA repair pathways (Lilley *et al*., 2011; Rodriguez *et al*., 2020; Smith & Weller, 2015; Weitzman *et al*, 2011) (Fig 8). New to this study, we demonstrate how host genome interactions are impacted by infection, revealing the loss of transcription regulatory factors and Pol II from cellular DNA, increased association with the nuclear pore, removal of the proteasome from host chromatin, and reorganization of host nucleoli. We also demonstrate that select DDR proteins interact with cellular DNA, while DNA repair proteins are enriched on viral genomes. This study not only provides a comprehensive view of nuclear reorganization during infection, but also establishes a novel approach to investigate how other DNA or RNA viruses convert host nuclei to hubs for viral production through manipulation of cellular processes.

**Figure 8.**
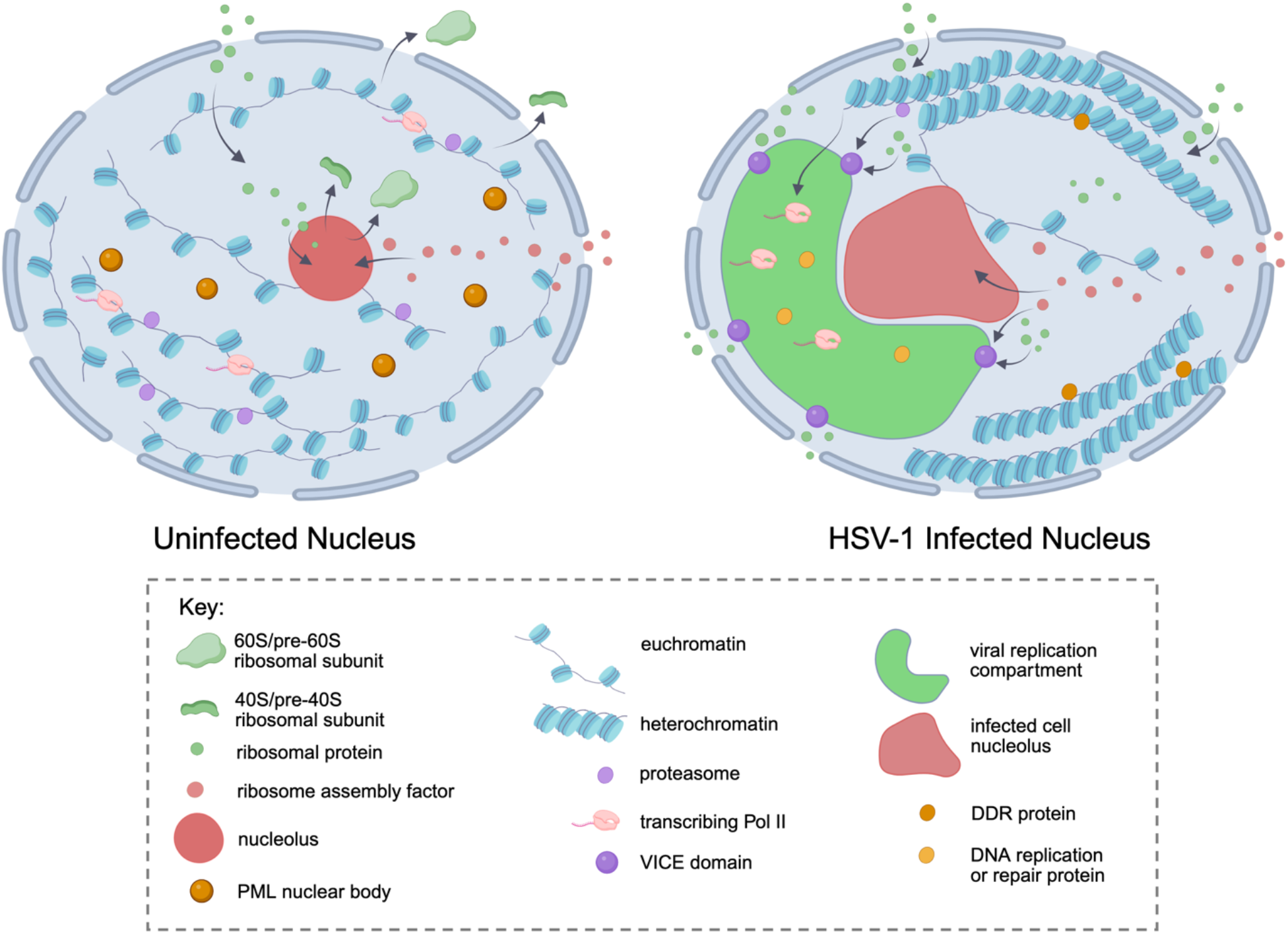
Model of HSV-1-mediated reorganization of nuclear proteins. Image generated in BioRender.

Comprehensive analysis of nuclear proteins has revealed five distinct groups of proteins that are reorganized during infection. These include proteins that are enriched on the viral genome, proteins that increase or decrease in association with the cellular genome, and proteins that increase or decrease in the nucleus. Below, we discuss each group.

Several proteins were identified to be enriched on viral DNA during infection. Select proteins appear to be redistributed from host chromatin to viral DNA, including chromatin remodeling factors, Pol II, the Mediator complex, TFIID, the Integrator complex, and mRNA processing factors. The levels of these factors do not change in the nucleus, and their association with cellular DNA generally decreases. These proteins are largely involved in active gene expression, consistent with preferential expression of viral genes during infection and shutdown of cellular Pol II-mediated transcription (Spencer *et al*, 1997). ICP4 recruits Mediator, TFIID, and Pol II to viral genes, which lack cellular chromatin, enabling this shift (Dremel & DeLuca, 2019). Additional proteins were found to associate with viral replication compartments (Fig 2), though no change in host genome association was detected. Within this group, PCNA and minichromosome maintence (MCM) protein levels increase in the nucleus, though nuclear levels of MMR and specific chromatin remodeling factors do not change. It is possible that early during infection, disruption of PML nuclear bodies by ICP0 releases these proteins from their subnuclear compartment, enabling association with viral DNA.

Shifting focus, select viral and DDR proteins, ribosome assembly factors, the nuclear pore, and the nuclear matrix increase in association with cellular DNA during infection. The increased association of viral proteins with cellular DNA is likely a reflection of their robust expression and nuclear localization by 6 hpi. Conversely, the relative nuclear levels of DDR proteins such as MDC1, most ribosome assembly factors, nuclear pore and nuclear matrix proteins do not change. Increased association with the nuclear pore may contribute to the mechanism by which host chromatin is marginalized to the periphery of the nucleus.

Because the nuclear levels of most ribosome assembly factors do not change during infection, it is possible that stalled ribosome biogenesis leads to the accumulation of assembly factors associated with cellular DNA (Belin *et al*., 2010). The factors identified largely include proteins involved in pre-60S processing in the nucleolus before release from rDNA (Broeck & Klinge, 2023). Although these factors copurify with host chromatin by aniPOND, that interaction is likely mediated through interactions with pre-rRNA tethered to the rDNA locus. We found that BRIX1 remains localized to the nucleolus during infection. However, the shape of the nucleolus shifts from a rounded structure to a less compact and structured compartment in most cells (Fig 7). This is consistent with infection stalling ribosome biogenesis (Belin *et al*., 2010; Metzger *et al*., 2025) and likely results in increased chromatin association of stalled preribosomes that fail to be released from the nucleolus. Others have demonstrated a distinct block in pre-60S assembly (Belin *et al*., 2010) and a decrease in the assembly of nascent ribosomes (Metzger *et al*., 2025) during infection. Data presented here provide mechanistic insight into the defects observed in pre-60S assembly. Small nucleolar ribonucleoprotein (snoRNP) associated proteins that modify pre-60S ribosomes and MDN1, which releases small snoRNPs and 60S assembly factors from pre-60S ribosomes, accumulate (Vanden Broeck & Klinge, 2024), demonstrating a distinct block in release from the nucleolus. Why this occurs rather than a complete shutdown of Pol I-mediated transcription of rRNA is not clear. However, we predict that this prevents a stress response (Szaflarski *et al*, 2022) or contributes to the increased fluidity of the nucleus during infection (Herzog *et al*., 2026).

We also observed distinct enrichment of DDR-related proteins histone H2AX, TP53BP1, and MDC1 on cellular DNA during infection. Although H2AX levels increase in the nucleus, TP53BP1 levels decrease, and MDC1 levels do not appear to change. These proteins play critical interconnected roles in the DDR following the formation of DNA double-strand breaks, as phosphorylation of H2AX leads to recruitment of MDC1, followed by TP53BP1 (Xie *et al*, 2007). These proteins form discrete foci at sites of damage, are activated by ATM phosphorylation, and contribute to amplification of ATM signaling, resulting in stalling of the cell cycle at the G1/S, intra-S, and G2/M checkpoints. ATM activation is proviral (Mertens & Knipe, 2021). Therefore, association of these proteins with cellular DNA may mediate a critical block in cell cycle progression to enable viral replication (Flemington, 2001), with downstream effectors blocked by ICP0-mediated degradation (Lilley *et al*., 2011).

Another subset of proteins is those that decrease association with cellular DNA during infection, including factors involved in active transcription, proteasome components, ICP0 target DNA-PK (PRKDC gene), and the chaperonin complex. Of this group, transcription factors shift localization to viral replication compartments as discussed above. The proteasome plays key roles in chromatin remodeling during active transcription and DNA replication (McCann & Tansey, 2014). As these processes are no longer active on cellular DNA during infection and viral DNA is not associated with chromatin, the proteasome is likely not needed for viral gene expression or on cellular DNA. It has been previously demonstrated that proteasome proteins accumulate in VICE domains during HSV-1 infection (Burch & Weller, 2004), consistent with their removal from host chromatin. The decrease in host chromatin association of ICP0 targets and the chaperonin complex is likely a reflection of decreased nuclear levels. Loss of DNA-PK helps to suppress non-homologous end joining during infection, as single-strand annealing is favored for productive infection (Lees-Miller *et al*, 1996; Smith & Weller, 2015).

Looking specifically at nuclear protein changes, ribosomal proteins and select histone proteins increase in the nucleus during infection. A recent study by the Cristea laboratory demonstrated that increased nuclear and VICE domain accumulation of ribosomal proteins likely contributes to disruption of ribosome biogenesis (Metzger *et al*., 2025). Why these proteins accumulate in VICE domains is unclear, but the intrinsically disordered tails of most ribosomal proteins may enable phase separation into this nuclear subcompartment (Peng *et al*, 2014). Although levels of some histone proteins increase in the nucleus, we did not observe increased association with cellular DNA. Because most histone proteins are synthesized during S phase (Nelson *et al*, 2002), it is not surprising that the levels of many histone proteins do not change. During chromatin compaction, the number of histones on cellular DNA does not increase, but nucleosomes are structurally reorganized closer together (Luger *et al*, 2012), which likely occurs during viral infection.

Finally, a set of proteins that decrease in the nucleus without a significant change in association with cellular DNA includes core components of PML nuclear bodies (PML, SP100, and SUMO1) and proteins involved in stress responses and mRNA processing. ICP0 targets PML proteins for proteasomal degradation (Rodriguez *et al*., 2020) and the spliceosome is disrupted by infection as most viral genes are not spliced (Hardy & Sandri-Goldin, 1994).

Taken together, we introduce a modification of aniPOND that enables the capturing of cellular genome-associated proteins during active viral DNA replication. When paired with nuclear proteomics and viral aniPOND, findings reveal a comprehensive view of nuclear restructuring during infection. This study provides new insight into viral modification of nuclei during HSV-1 infection and highlights the utility of this approach to investigate infection with less well studied DNA and even RNA viruses. This approach can also be used to determine how different cell types respond to infection and how nuclei are remodeled during infection with viral mutants and different stages of infection, including latency and reactivation.

## Materials and Methods

### Cells and Viruses

Experiments were conducted using MRC-5 cells (human embryonic lung, CCL-171) obtained from and propagated as directed by ATCC. HSV-1 strain KOS (Neal DeLuca) was used for all infections.

### CellTiter-Glo Luminescent Cell Viability Assay

CellTiter-Glo Luminescent Cell Viability Assay (Promega, G7571) was carried out to determine the effect of EdC (5-Ethynyl-2′-deoxycytidine, Sigma, T511307) on cell viability. EdC was dissolved in DMSO to a stock concentration of 40 mM. MRC-5 cells were plated at a density of 1.25×10^4^ cells/well in a 96-well plate. After 24 hours, EdC was diluted and added to the cells at concentrations indicated in Figure S1. Cells were treated with EdC for 48 hours before conducting the CellTiter-Glo assay according to the manufacturer’s protocol. Luminescence was measured on a SpectraMax iD3 plate reader. Percent viability was measured as the relative light units (RLU) of each EdC-treated condition divided by the control group of cells treated with 1% DMSO.

### Immunofluorescence and Imaging of EdC-Labeled DNA

MRC-5 cells were plated on coverslips in a 12-well dish at a density of 1×10^5^ cells/well. 24 hours after plating, 25 μM EdC was added to the cell culture medium to label replicating cellular DNA. After 48-hours of incubation, EdC was washed off, and cells were incubated for an additional 24 hours. Cells were then infected with HSV-1 strain KOS at an MOI of 10 PFU/cell, and infection was carried out for 6 hours. For comparison, a mock-infected control was included. To label viral and not cellular DNA, cells were incubated without EdC for 72 hours, then 25 μM EdC was added to the cell culture medium from 4 to 6 hpi. Cells were fixed with 4% paraformaldehyde for 15 minutes at room temperature, and coverslips were incubated in 1x phosphate-buffered saline (PBS) with 3% bovine serum albumin (BSA) overnight. Cells were then permeabilized with 0.5% Triton X-100 in 1x PBS for 20 minutes and blocked with 3% BSA in 1x PBS for 30 minutes. EdC-labeled DNA was then covalently linked to Alexa Fluor 488 Azide (Invitrogen, A-10266) for 30 minutes using a click chemistry reaction (Dembowski & Deluca, 2017). Cells were then stained with 300 nM DAPI for 30 minutes, followed by another 30-minute incubation of primary antibodies mouse α-ICP4 (Abcam, AB6514, 1:200) or rabbit α-BRIX1 (Sigma, HPA039614, 1μg/mL) diluted in 1x PBS. Cells were washed with 1% BSA in 1x PBS, then 1x PBS three times. Finally, cells were incubated with the secondary antibody, Alexa Fluor 594-conjugated goat α-mouse (Invitrogen, A11005, 1:200) or Alexa Fluor 594-conjugated goat α-rabbit (Invitrogen, A11012, 1:200) in 1x PBS for 30 minutes. Cells were washed with 1x PBS, mounted onto glass slides with Shandon Immu-Mount, and imaged using a Nikon Eclipse Ti2 Inverted Confocal Microscope.

### aniPOND

aniPOND was conducted as described previously (Dembowski & Deluca, 2017; Leung *et al*., 2013). MRC-5 cells were plated in 600 cm^2^ dishes (1.2×10^6^ cells/plate, ∼20% confluent). After 24 hours, 25 μM EdC was added to the cell culture medium to label replicating cellular DNA. After 48 hours, EdC was washed off, and cells were incubated for an additional 24 hours. Cells were then either mock infected or infected with KOS at an MOI of 10 PFU/cell, followed by a 6-hour incubation (Figure 1A). To label viral and not cellular DNA, cells were plated and grown under the same conditions without EdC, infected, and then 25 μM EdC was added to the cell culture medium from 4 to 6 hpi. We previously confirmed that under these conditions viral DNA and not cellular DNA is labeled with EdC (Dembowski & DeLuca, 2015; Dembowski *et al*., 2017). Each condition also had a negative control without EdC. Nuclear extraction, click chemistry, cell lysis, and sonication were carried out as described previously (Dembowski & Deluca, 2017, 2018), except cell lysis occurred for 30 minutes on ice followed by sonication 8 times at 40% amplitude with a 3 mm probe. Proteins were eluted from streptavidin-coated beads by boiling in 2X SDS Laemmli sample buffer.

### Nuclear Protein Isolation

Cells were plated and infected or mock infected according to the aniPOND protocol above. Nuclear proteins were isolated following the nuclei lysis step of aniPOND. The click reaction was prepared and added to the cells without biotin azide. Following nuclear lysis, proteins were TCA precipitated and resuspended in 2X SDS Laemmli sample buffer.

### Mass Spectrometry and Data Analysis

MS was carried out by MSBioworks as previously described (Dembowski & DeLuca, 2015). Proteins were determined enriched by aniPOND on certain populations of DNA based on the following criteria and analyses. To account for background, proteins were removed if they did not have at least 4 spectral counts in two (or more) replicates of any positive condition. Spectral abundance factor (SAF) was calculated for each protein by dividing the number of spectral counts by the protein’s molecular weight, thus accounting for variation in protein size. Next, the fold enrichment of the SAF was calculated by dividing the SAF of a positive condition by the corresponding negative control (no EdC). To further eliminate background proteins, only those with at least 2-fold SAF enrichment in (at least) 2 replicates of any comparison were kept for further analysis. Next, the NSAF was calculated by dividing the SAF by the total number of spectral counts for the sample. Averaged NSAF values from 3 biological replicate data sets were used for further analysis.

Proteins were considered enriched in the nuclear preparation if they had at least 4 spectral counts in 2 or more replicates of either condition (infected or mock). NSAF was calculated as explained above to account for variation in protein size and relative abundance. Average NSAF values from 3 biological replicate data sets were used for downstream analysis.

Proteins enriched on infected cell DNA were compared to proteins enriched on viral DNA, uninfected cell DNA, and nuclear proteins, as described in the results. Comparisons were used to calculate p-value and fold change. Volcano plots were generated in GraphPad Prism, and Venn diagrams were generated using BioTools.fr. STRING interaction networks were created using string-db.org (Szklarczyk *et al*, 2019). A full STRING network type with confidence edges was used, and sources of interactions included text mining, experiment, databases, co-expression, neighborhood, gene fusion, and co-occurrence. The confidence level was set to high (0.700) and disconnected nodes were hidden but included in supplemental material (Fig S1, Table S1, Table S2) or Venn Diagrams (Figs. 3 and 5). Markov’s Clustering Algorithm was used with an inflation parameter of 3, and the edges between clusters were not shown.

### Analysis of Circularity of the Nucleolus

Immunofluorescence and imaging of EdC-labeled DNA was completed as stated earlier. BRIX1 protein was used as a marker for the nucleolus. Images were analyzed using imageJ (version 1.54p/Fiji). Prior to quantification, images were thresholded using color threshold to highlight the shape against the background. The scale was set using the set scale function in the software after measuring the scale bar present in the image. The nucleolus was traced in the TRITC channel using the freehand tool. After tracing the complete shape, the area, perimeter and circularity were measured using the measure tool. Circularity values range from 0.0 (infinitely elongated polygon) to 1.0 (perfect circle).

## Acknowledgements

We would like to thank members of the Dembowski Lab for their helpful feedback and discussions. We would also like to thank Dr. Neal DeLuca (University of Pittsburgh) for HSV-1 strain KOS and Rachel Lloyd (University of California, Berkeley) for help with statistical analyses. The confocal microscope used in this study was purchased using funds provided to Dr. John Pollock (Duquesne University) from the National Science Foundation (DBI Award #1726368).

## Supporting Information

**Figure S1. Effect of EdC labeling on cell viability.** A CellTiter-Glo Luminescent cell viability assay was performed to assess the cytotoxic effects of treating cells with EdC at the indicated concentrations for 48 hours. The mean viability of cells treated with EdC was compared with that of the negative control, cells treated with 1% DMSO (equivalent to the volume of DMSO in the highest concentration of dissolved EdC). Each data point represents the mean of three biological replicates. A one-way ANOVA with Dunnett’s multiple comparisons test was used to compare each EdC-treated condition to the negative control. * p ≤ 0.05.

**Figure S2. Disconnected nodes from STRING diagrams.** Proteins that were enriched on the viral (A) or cellular (B) genomes during infection but not included in the STRING diagrams in Figures 2B and 2C because predicted interactions fell below the interaction score (high confidence, 0.700) are listed.

**Table S1. Raw data, processing, and analysis summary for aniPOND mass spectrometry data.**

**Table S2. Raw data, processing, and analysis summary for total nuclear proteins.**

