## Supplemental Figures 1 & 2 for "Distinct labeling of host and viral genomes highlights redistribution of nuclear proteins during HSV-1 infection"

### Supporting Information

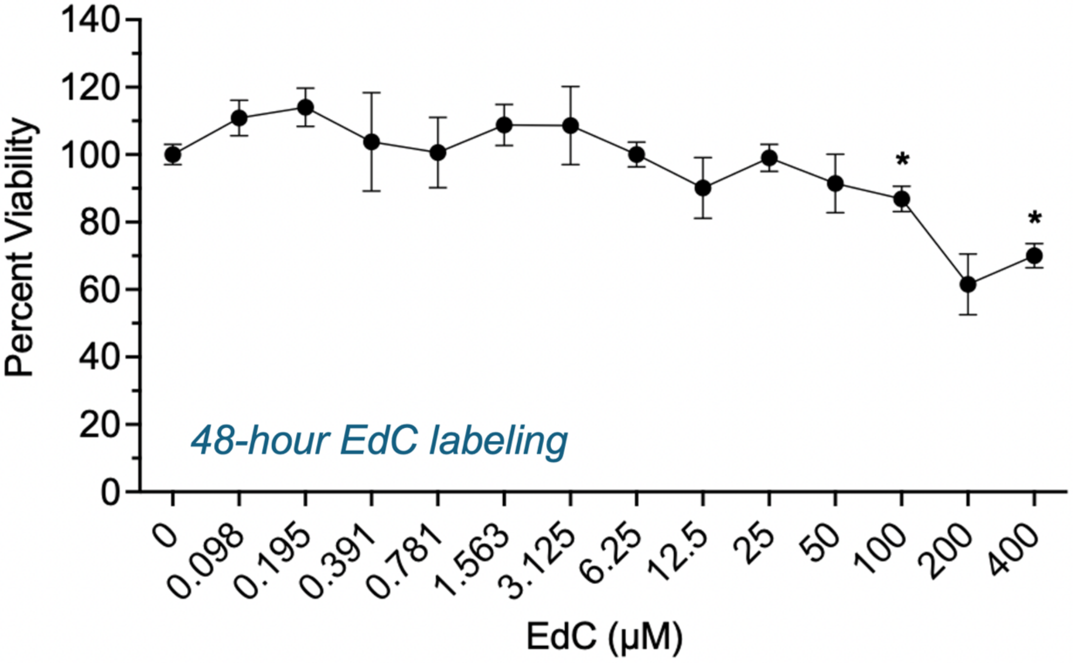


**Figure S1. Effect of EdC labeling on cell viability.** A CellTiter-Glo Luminescent cell viability assay was performed to assess the cytotoxic effects of treating cells with EdC at the indicated concentrations for 48 hours. The mean viability of cells treated with EdC was compared with that of the negative control, cells treated with 1% DMSO (equivalent to the volume of DMSO in the highest concentration of dissolved EdC). Each data point represents the mean of three biological replicates. A one-way ANOVA with Dunnett's multiple comparisons test was used to compare each EdC-treated condition to the negative control. * $p \leq 0.05$.

A

*Enriched on viral genome at 6 hpi*
*Disconnected nodes:*

| | |
|---|---|
| ANPEP | MTHFD1L |
| CAV1 | MYOF |
| CNN3 | NFIC |
| EMILIN1 | PLAT |
| EPB41L2 | PLOD3 |
| ERLIN2 | PSMB6 |
| FARP1 | SAMD1 |
| GLG1 | SMCHD1 |
| GPC1 | SNTB2 |
| HNRNPH3 | TGM2 |
| IFI16 | |

B

*Enriched on host genome at 6 hpi*
*Disconnected nodes:*

| | |
|---|---|
| ARPC5 | MOGS |
| ATP6V0A1 | MYEF2 |
| BAZ2A | NEU1 |
| CARD19 | POGZ |
| CCAR2 | PPP1R18 |
| CIRBP | PRDX6 |
| CKAP2 | PRPH |
| DDB1 | PSPC1 |
| DYNC1H1 | RAI14 |
| GDF15 | RBM12B |
| GREM1 | RBM3 |
| HIST1H10 | RBMXL1 |
| HIST1H11 | SLTM |
| KRT18 | SUB1 |
| KTN1 | SVIL |
| LACTB | TRIOBP |
| LARP4 | TRPV2 |
| LEMD2 | YLPM1 |
| LIMA1 | ZC3HAV1 |
| MACROH2A2 | ZNF638 |
| MAP4 | |

**Figure S2. Disconnected nodes from STRING diagrams.** Proteins that were enriched on the viral (A) or cellular (B) genomes during infection but not included in the STRING diagrams in Figures 2B and 2C because predicted interactions fell below the interaction score (high confidence, 0.700) are listed.

**Table S1. Raw data, processing, and analysis summary for aniPOND mass spectrometry data.**

**Table S2. Raw data, processing, and analysis summary for total nuclear proteins.**
